# Bayesian Optimization for Ternary Complex Prediction (BOTCP)

**DOI:** 10.1101/2022.06.03.494737

**Authors:** Arjun Rao, Tin M. Tunjic, Michael Brunsteiner, Michael Müller, Hosein Fooladi, Noah Weber

## Abstract

Proximity-inducing compounds (PICs) are an emergent drug technology through which a protein of interest (POI), often a drug target, is brought into the vicinity of a second protein which modifies the POI’s function, abundance or localisation, giving rise to a therapeutic effect. One of the best-known examples for such compounds are heterobifunctional molecules known as proteolysis targeting chimeras (PROTACs). PROTACs reduce the abundance of the target protein by establishing proximity to an E3 ligase which targets the protein towards degradation via the ubiquitin-proteasomal pathway. Design of PROTACs in silico requires the computational prediction of the ternary complex consisting of POI, PROTAC molecule, and the E3 ligase.

Here, we present a novel machine learning-based method for predicting PROTAC-mediated ternary complex structures using Bayesian optimization. We show how a fitness score combining an estimation of protein-protein interactions with PROTAC binding energy calculations enables the sample-efficient exploration of candidate structures. Furthermore, our method presents two novel scores for filtering and reranking which take PROTAC stability (Autodock-Vina based PROTAC stability score) and protein interaction restraints (the TCP-AIR score) into account. We evaluate our method using DockQ scores and demonstrate, that even with a clustering that require members to have a high similarity, i.e. with smaller clusters, we can assign high ranks to those clusters that contain poses close to the experimentally determined native structure of the ternary complexes. We also demonstrate the resultant improved yeild of near-native poses in these clusters.

## 1 Introduction

Targeted protein degradation is an emerging therapeutic modality which, instead of inhibiting the activity of a drug target, acts by inducing degradation of the target protein itself [1, 2]. It employs so-called monofunctional degraders such as molecular glues or heterobifunctional degraders such as Proteolysis Targeting Chimeras (PROTACs) which act as proximity-inducing compounds. PICs bring together the protein of interest (POI) and an E3 ubiquitin ligase to induce subsequent ubiquitination and degradation of the target protein (See Figure 1) [3, 4, 5, 6].

**Figure 1.**
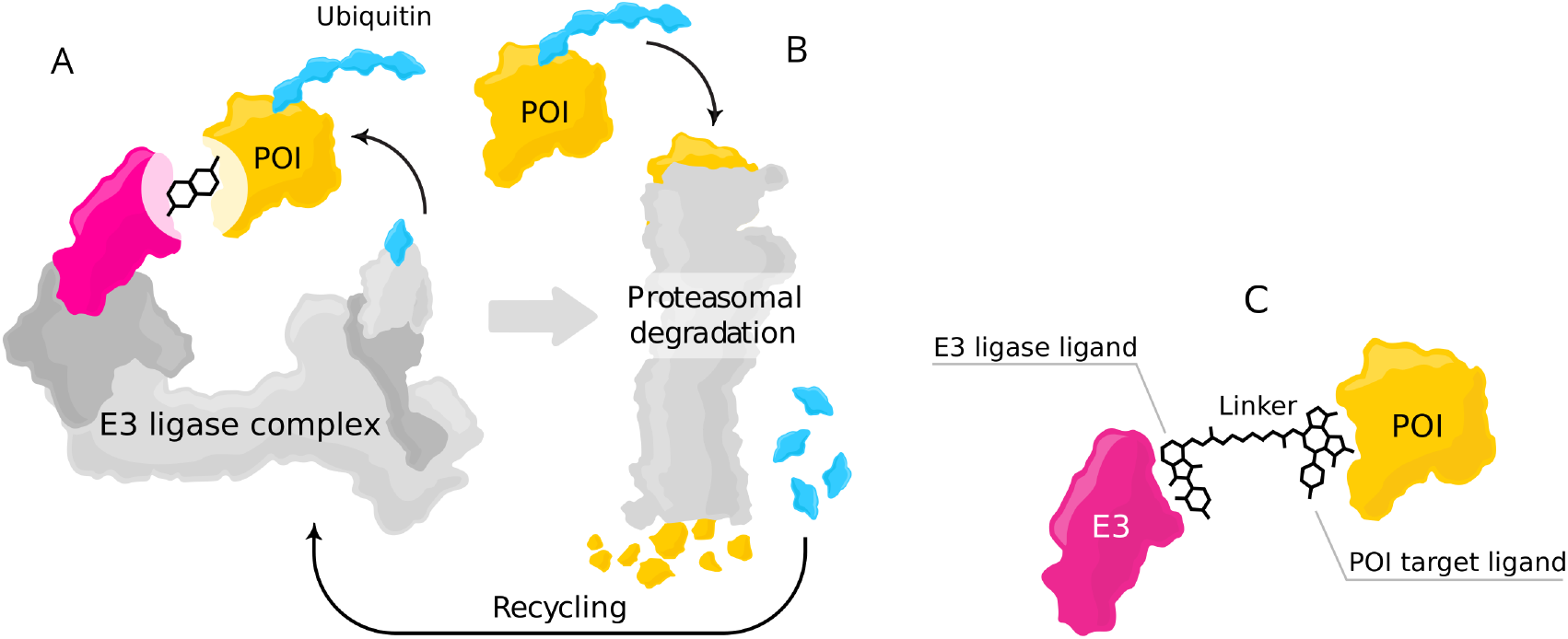
Mode of action for targeted protein degradation: (A) A POI is shown in a proximity with an E3 ligase complex – the receptor part of E3 ligase is highlighted in magenta – a heterobifunctional degrader such as PROTAC brings together the POI and E3 ligase, which initiate the degradation process. (B) After formation of a ternary complex, the POI is marked with a polyubiquitin chain (blue), which subsequently leads to protein degradation through UPS. After degradation, the ubiquitin molecules are detached and recycled by the UPS. Moreover, the PROTAC molecule is detached and can induce degradation of multiple POIs (C) Ternary complex formed from the interaction of the POI, receptor part of the E3 ligase and heterobifunctional degrader.

Targeted protein degradation using PROTACs offers several advantages over traditional occupancy-based inhibitors such as [7]:

- Expanding the druggable space: there is no need for tight binding to the functional site of the POI, and as a result, PROTACs can target proteins more shallow binding pockets (e.g., transcription factors and scaffolding proteins).
- Working at a substoichiometric concentration: A single PROTAC molecule can induce degradation of multiple POI molecules over time.
- Improved efficacy: Recovery of target protein abundance (and therefore overall activity) is linked to protein synthesis in cells and not just to the absence of the PROTAC.
- Improved selectivity: PROTACs are typically larger than traditional inhibitors and can utilize more binding sites on the target which results in higher selectivity.

Structurally, a PROTAC consists of a warhead that binds to the POI, an E3 ligand that binds into the E3 ligase, and a linker that connects these two parts. Current PROTACs utilize a limited number of E3 ligases. The most prominent E3 ligases are cereblon (CRBN) and Von-Hippel-Lindau Tumor Suppressor (VHL) due to the availability of high-affinity binders and favourable properties for both proteins. For CRBN binders, immunomodulatory drugs such as thalidomide [8], lenalidomide [9], and pomalidomide [10] are widely used. For VHL, binders were initially derived from a natural degron peptide and evolved to highly potent small molecules [11, 12]. In addition, some other E3 ligases such as cell inhibitors of apoptosis protein (cIAP) [13], murine double minute 2 (MDM2) [14], and KEAP1 [15] have been harnessed for targeted protein degradation. There is an ongoing effort on expanding the library of E3 ligases and finding suitable potent ligands against other E3 ligases.

While the set of E3 ligases used for targeted protein degradation with PROTACs has so far been limited, they have been used to target a diverse range of POIs from different protein families. There are prominent examples with good evidence of safety and efficacy in initial clinical trials, including degraders targeting the androgen receptor (AR) [16, 17], estrogen receptor (ER) [18], IRAK4, BCL-XL [19], Helios (IKZF2) [20], and GSPT1. Furthermore, in vivo studies for PROTAC targeting BRD4 [21], BTK [22, 23], RIPK2 [24], and SMARCA2 [25] have shown promising results. These findings highlight the potential of PROTAC for the degradation of diverse protein families.

Judging the efficacy of a candidate PROTAC molecule in-silico is a daunting task owing to a plethora of interactions that are difficult to evaluate. For a PROTAC to be effictive, it must at-least satisfy the following requirements. Firstly, it must be able to penetrate the cell membrane and stay stable inside the cell. Secondly, its interactions with the E3 ligase and the POI must result in the formation of a stable ternary complex where the POI is held near the E3 ligase. Finally, this complex must be one that lends itself to being tagged by ubiquitin so that the POI may be degraded.

In light of the above steps, it is clear that accurate and efficient prediction of the structure of the ternary complex is imperative for improvements at many stages of the PROTAC development process. For instance, computational modeling of ternary complexes allows the screening of multiple different E3 ligases for a candidate target, maximising the stability of the ternary complex [26]. Knowing the structure of the ternary complex is the first step to calculating its stability, and experimental evidence shows that the formation of a stable ternary complex can lead to more efficient degradation [26].

However ternary complex prediction is in itself a challenging task as one needs to take the interactions between several components into account. For example, the interaction between the warhead and the E3-binding fragments and their respective proteins, the protein-protein interaction between the E3-ligase and the target protein, as well as the interactions and physical constraints imposed by the linker. Furthermore it requires the ability to score the validity of any single physical configuration of the two proteins and the degrader, and to optimize this score to ultimately predict the structure of the ternary complex efficiently. Even if we know the fragments that bind to the E3 ligase and the POI, what constitutes an optimal linker is not yet fully understood [27]. The linker length and composition can have a significant impact on cooperativity and stability of the ternary complex [28, 22], and consequently on the efficiency of the degradation [29].

Accurate and efficient ternary complex prediction is imperative for improvements at many stages of the PRO- TAC development process. For instance, computational modeling of ternary complexes allows the screening of multiple different E3 ligases for a candidate target, maximising the stability of the ternary complex [26]. It can further facilitate linker design by investigating the impact of the linker on the ternary complex. These steps can incorporate different warheads, finding an ideal combination with available E3 ligase and binders and a linker.

In one of the first attempts for computationally predicting ternary complexes, Drummond et al. [30] proposed four different methods for the prediction ensemble of PROTAC mediated ternary complexes. With the most promising method of combining docking of the two protein-ligand structures with separate PROTAC conformer generation, they showed that it is possible to recover near-native ternary complexes for a few available PDBs. In follow-up work [31], Drummond et al. added two additional methods which improve the previous performance. This work introduces clustering of the generated structures, an essential step in obtaining good results. Zaidman et al. [32] proposed another method, PRosettaC, which adds consecutive steps such as global protein-protein docking and local docking refinement, generating conformation of PROTACs that can fit in the docking solutions and final clustering. Although they have been able to reproduce a good amount of crystal structures of ternary complexes, they are initiating their prediction from bound structures, which does not reflect real-world conditions. Bound structures come from the decomposition of already available ternary complexes, but *a priori* we do not have access to this information. In one of the recent works, Weng et al. [33] proposed a method again based on global protein-protein docking, local refinement with RosettaDock [34], generating PROTACs using RDkit, and applying filtering and re-ranking criteria for predicting near-native ternary complexes. They started from unbound structures and showed that good results could be achieved with these methods.

The first contribution of our current work, is the demonstration of the successful use of a Bayesian optimization (BO) [35] framework to perform a sample-efficient exploration of the space of ternary complex candidates. BO is an established method which allows optimizing functions which are costly to evaluate by proposing promising evaluation candidates (here: candidate ternary complex structures). This leads to sample efficiency, i.e., fewer candidates need to be evaluated (e.g. by a docking software) compared to (e.g. Monte Carlo-) search methods, and allows the use of potentially complex fitness functions which are needed to characterize the multiple interactions within a ternary complex. We use this to sample poses scored by a weighted sum of a protein-protein interaction (PPI) score and a PROTAC score to model the feasibility of a candidate ternary complex structure. The PPI score quantifies the interactions between the proteins in a particular candidate, while the PROTAC score takes into account the feasibility of embedding a conformation of the PROTAC molecule given the position of the binding pockets in this configuration of proteins.

The second contribution of our work lies in the design of two novel scores specialized for filtering, ranking and clustering. The first score is used for filtering favourable complexes - a version of the ambiguous interaction restraints energy (here abbreviated as “TCP-AIR Energy”) [36] that has been modified specifically for ternary complexes and prioritizes poses with ideal protein configurations. The second score is the PROTAC stability score, which is calculated using a modified version of Autodock-Vina. It packs the PROTAC and indicates its stability given a particular configuration of proteins. This score is useful for ranking.

We show in our current work (See table 2) that, with the efficient sampling with the BO loop, combined with filtering, clustering, and ranking using the above scores, we are able to effectively find highly ranked clusters with near-native poses. Specifically, compared to previous works, our clustering is more confined and each cluster contains only a small number of points. Our ability to effectively rank these, gives us clusters which have a high yield of near-native scores.

**Table 1:** Details of the protein structures used in this work. Columns show the PDB ID for each ternary complex as well as the chain identifiers (complex), the gene name, the PDB ID of the structure used as input for prediction (template), the chain IDs of E3 ligase and POI, the first and last residue number included into the used models (N-term, C-term). For the PROTAC, the colums show the residue ID, the nature of any ionizable group, and the net charge used in molecular models.

| complex | E3 ligase |  |  |  |  | POI |  |  |  | PROTAC |  |  |  |
| --- | --- | --- | --- | --- | --- | --- | --- | --- | --- | --- | --- | --- | --- |
|  | Name | Template | Chain | N-term | C-term | Name | Template | Chain | N-term | C-term | Residue | Ionizable | Net charge |
| 5T35-DA | VHL | 4W9H-I | D | 62 | 204 | BRD4-2 | 5UEU-A | A | 349 | 457 | 759 | no | 0 |
| 5T35-HE | VHL | 4W9H-I | H | 62 | 204 | BRD4-2 | 5UEU-A | E | 349 | 457 | 759 | no | 0 |
| 6BN7-bc | CRBN | 4TZ4-C | B | 48 | 426 | BRD4-1 | 3MXF-A | C | 44 | 168 | RN3 | imine | 0 |
| 6BOY-bc | CRBN | 4TZ4-C | B | 48 | 426 | BRD4-1 | 3MXF-A | C | 44 | 168 | RN6 | imine | 0 |
| 6HAX-BA | VHL | 4W9H-I | B | 62 | 204 | SMARCA2 | 6HAZ-A | A | 1378 | 1490 | FWZ | piperazine | 1 |
| 6HAX-FE | VHL | 4W9H-I | F | 62 | 204 | SMARCA2 | 6HAZ-A | E | 1378 | 1490 | FWZ | piperazine | 1 |
| 6HAY-BA | VHL | 4W9H-I | B | 62 | 204 | SMARCA2 | 6HAZ-A | A | 1378 | 1490 | FX8 | piperazine | 1 |
| 6HAY-FE | VHL | 4W9H-I | F | 62 | 204 | SMARCA2 | 6HAZ-A | E | 1378 | 1490 | FX8 | piperazine | 1 |
| 6HR2-BA | VHL | 4W9H-I | B | 62 | 204 | SMARCA4 | 6ZS2-A | A | 1449 | 1569 | FWZ | piperazine | 1 |
| 6HR2-FE | VHL | 4W9H-I | F | 62 | 204 | SMARCA4 | 6ZS2-A | E | 1449 | 1569 | FWZ | piperazine | 1 |
| 6SIS-DA | VHL | 4W9H-I | D | 62 | 204 | BRD4-2 | 5UEU-A | A | 349 | 457 | LFE | sec amine | 1 |
| 6SIS-HE | VHL | 4W9H-I | H | 62 | 204 | BRD4-2 | 5UEU-A | E | 349 | 457 | LFE | sec amine | 1 |
| 6W7O-CA | BIRC2 | 6W74-A | C | 266 | 349 | BTK | 5P9J-A | A | 396 | 656 | TL7 | sec amine | 1 |
| 6W7O-DB | BIRC2 | 6W74-A | D | 266 | 349 | BTK | 5P9J-A | B | 396 | 656 | TL7 | sec amine | 1 |
| 6W8I-DA | BIRC2 | 6W74-A | D | 266 | 349 | BTK | 5P9J-A | A | 396 | 656 | TKY | sec amine | 1 |
| 6W8I-EB | BIRC2 | 6W74-A | E | 266 | 349 | BTK | 5P9J-A | B | 396 | 656 | TKY | sec amine | 1 |
| 6W8I-FC | BIRC2 | 6W74-A | F | 266 | 349 | BTK | 5P9J-A | C | 396 | 656 | TKY | sec amine | 1 |
| 6ZHC-ad | VHL | 4W9H-I | A | 62 | 204 | BCL2L1 | 4QVX-A | D | 2 | 197 | QL8 | carboxylic acid | -1 |
| 7JTO-LB | VHL | 4W9H-I | L | 62 | 204 | WDR5 | 4QL1-A | B | 32 | 333 | VKA | 2 piperazines | 2 |
| 7JTP-LA | VHL | 4W9H-I | L | 62 | 204 | WDR5 | 4QL1-A | A | 32 | 333 | X6M | piperazine | 1 |
| 7KHH-CD | VHL | 4W9H-I | C | 62 | 204 | BRD4-1 | 3MXF-A | D | 44 | 168 | WEP | no | 0 |
| 7Q2J-CD | VHL | 4W9H-I | C | 62 | 204 | WDR5 | 4QL1-A | D | 32 | 333 | 8KH | piperazine | 1 |

**Table 2:**
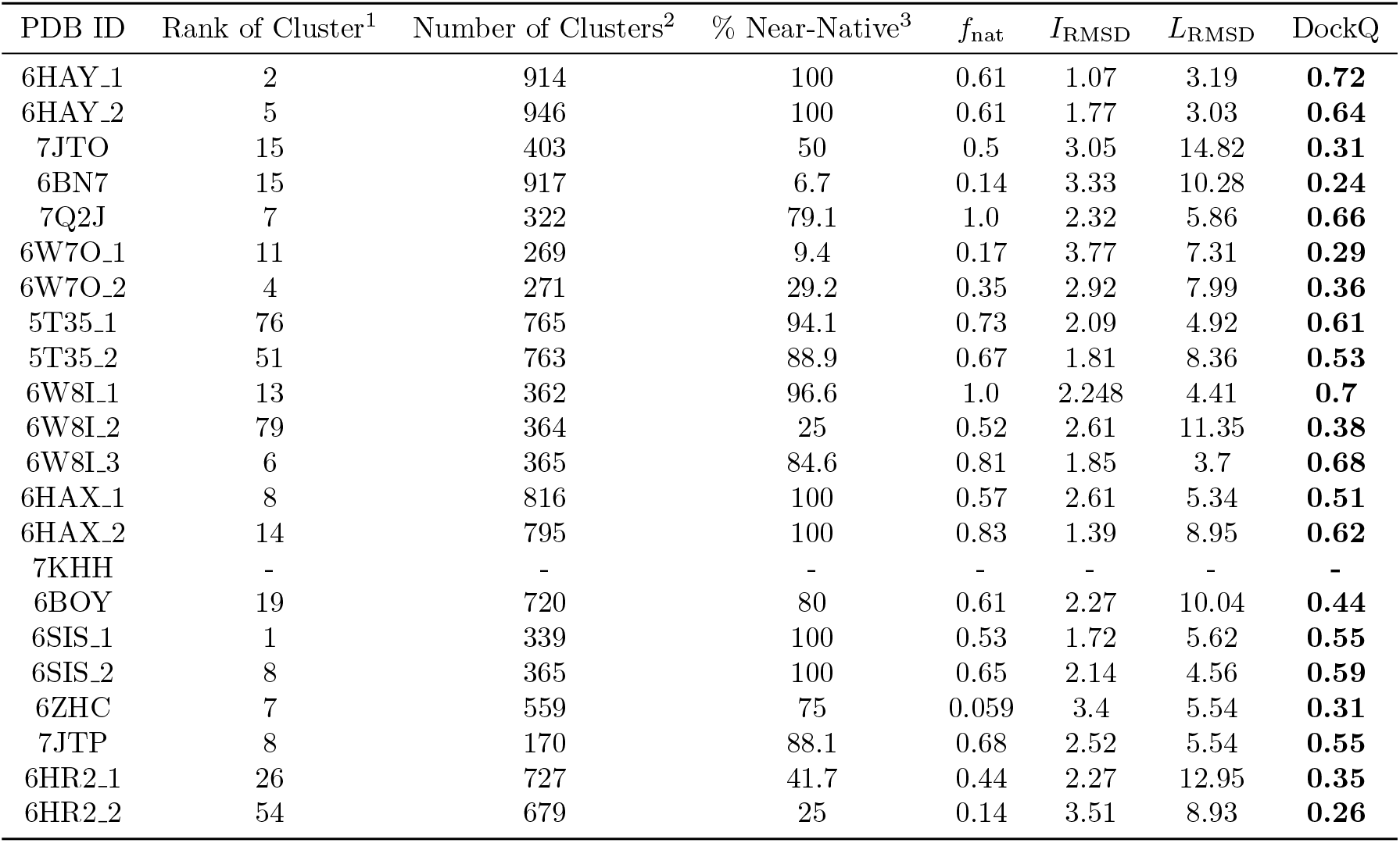
Results of the BOTCP method for ternary complex prediction on unbound structures before the refinement. ^1^The best rank containing at least one model with DockQ≥0.23. ^2^The total number of clusters. ^3^The percentage of the near-native poses in that specific ranked cluster. For each ternary complex, we selected a cluster which includes an RRT with a high DockQ score. Columns also show the PDB ID and characterization of the best RRT in terms of *f*_nat_, *I*_RMSD_, *L*_RMSD_ and DockQ score (see text).

## 2 Methods

In this paper, we propose the Bayesian Optimization for Ternary Complex Prediction (BOTCP) method. At its core lies a Bayesian optimization (BO) loop capable of finding appropriate ensembles of ternary complexes by optimizing an objective function which includes a fitness describing the strength of the interaction of the two proteins (“PPI fitness”) and a fitness modeling the possibility of connecting the binding pockets with the PROTAC (“PROTAC fitness”). We then perform a local optimization around the most promising candidates using a score which takes into account the stability of the PROTAC between the two proteins (“PROTAC stability score”, calculated by our own extension to Autodock-Vina). Afterwards, we cluster the resulting candidates before performing filtering using an extension to the ambiguous interaction restraint (AIR) energy [36], and re-ranking the final selections using the PROTAC stability score. In a nutshell, BOTCP consists of the following steps/stages:

1. Input initialization (Section 2.1)
2. Bayesian optimization (Section 2.2)
3. Local optimization with simulated annealing of the PROTAC stability score (Section 2.3)
4. Clustering, filtering using TCP-AIR energy, and re-ranking (Section 2.4)
5. Refinement (Section 2.5)

### 2.1 Initialization

At the time of writing, the protein database (PDB [37]) contained fifteen experimentally (x-ray) determined ternary complex structures which include a PROTAC molecule, a protein of interest, and an E3 ligase. We discarded one of these (7PI4) since here the POI is a kinase with a large flexible loop that is close to the proteinprotein interface and the PROTAC molecule but not resolved in the experimental structure. The remaining 14 structures are listed in Table 2.1. Each structure was downloaded from the PDB in its native format and preprocessed as follows. All (non-PROTAC) hetero atoms were removed, and the chains corresponding to the POI and the E3 ligase as well as the PROTAC molecule were saved in a single PDB file for comparison with predictions. In several cases, the asymmetric unit contained two or more ternary complexes, which were saved individually. The protein structures used as input for modeling were taken from other PDB entries (unbound structures) so as to emulate the practical use case where the experimental structure of the ternary complex is not available. The structures used here include three different E3 ligases and seven POIs. For each of these ten proteins, a PDB entry was selected according to the following criteria:

1. The protein has a bound ligand that is similar to the warhead or binder of the corresponding PROTAC molecule.
2. The structure comes with few and/or short gaps/unresolved loops and the resolution (R-factor) is good.

For one structure (7KHH), no PDB entry fulfilling the first criterion was available. Here we estimated the structure of the bound warhead by docking the warhead into the binding pocket, using the GNINA docking tool [38] with default parameters. The quality of the resulting complex structure was ascertained by visual inspection and comparison to the corresponding portion of the ternary complex structure. The resulting warhead and binder molecules (only non-hydrogen atoms) were saved in SDF format for usage as inputs to the BO method. Each of the ten individual single structures of E3 ligase or POI were then preprocessed using the following steps:

1. All heteroatoms (including water and small molecules) were removed while keeping ions (in four cases, the structure contained a Zinc ion complexed by four cysteine or histidine residues).
2. For those structures that had gaps, we downloaded the corresponding AlphaFold structures [39, 40], aligned the experimental and the AlphaFold structures using Chimera [41], and used the coordinates of the AlphaFold structures to fill the gaps.
3. We cropped flexible N- and C-terminal tails (usually with poor confidence levels in AlphaFold predictions) as specified in 2.1.
4. Missing atoms were replaced using the PDBfixer tool [42].
5. We used Reduce[43] to protonate/add hydrogens, and, where appropriate, flip asparagine, glutamine, or histidine side chains.

The PROTAC molecules were extracted from the original PDB file and converted to SDF format using Open Babel [44]. If required, bond orders and protonation were corrected manually. Here we assumed that all weak bases (secondary and tertiary amines) and acids (carboxylic acid) are (de)protonated to form anions or cations. In a few cases, the presence of piperazines required a choice to protonate one of the two nitrogens. This choice was made based on chemical intuition and the corresponding expected pKA variations. All SDF files are provided in the SI. The protein PDB and PROTAC SDF files resulting from this procedure were used as input for the BO loop.

Our method uses three inputs: the POI with the docked warhead, the E3 ligase with the docked E3 binder, and the PROTAC SMILES (see Figure 2 for an example), which are provided to the Bayesian optimization loop. It is more suitable to access the co-crystal structure of POI+warhead and E3 ligase+binder, but if this information is not available, it is required first to dock the corresponding ligands into their proteins and use these structures after molecular docking for the next step in out BOTCP pipeline.

**Figure 2:**
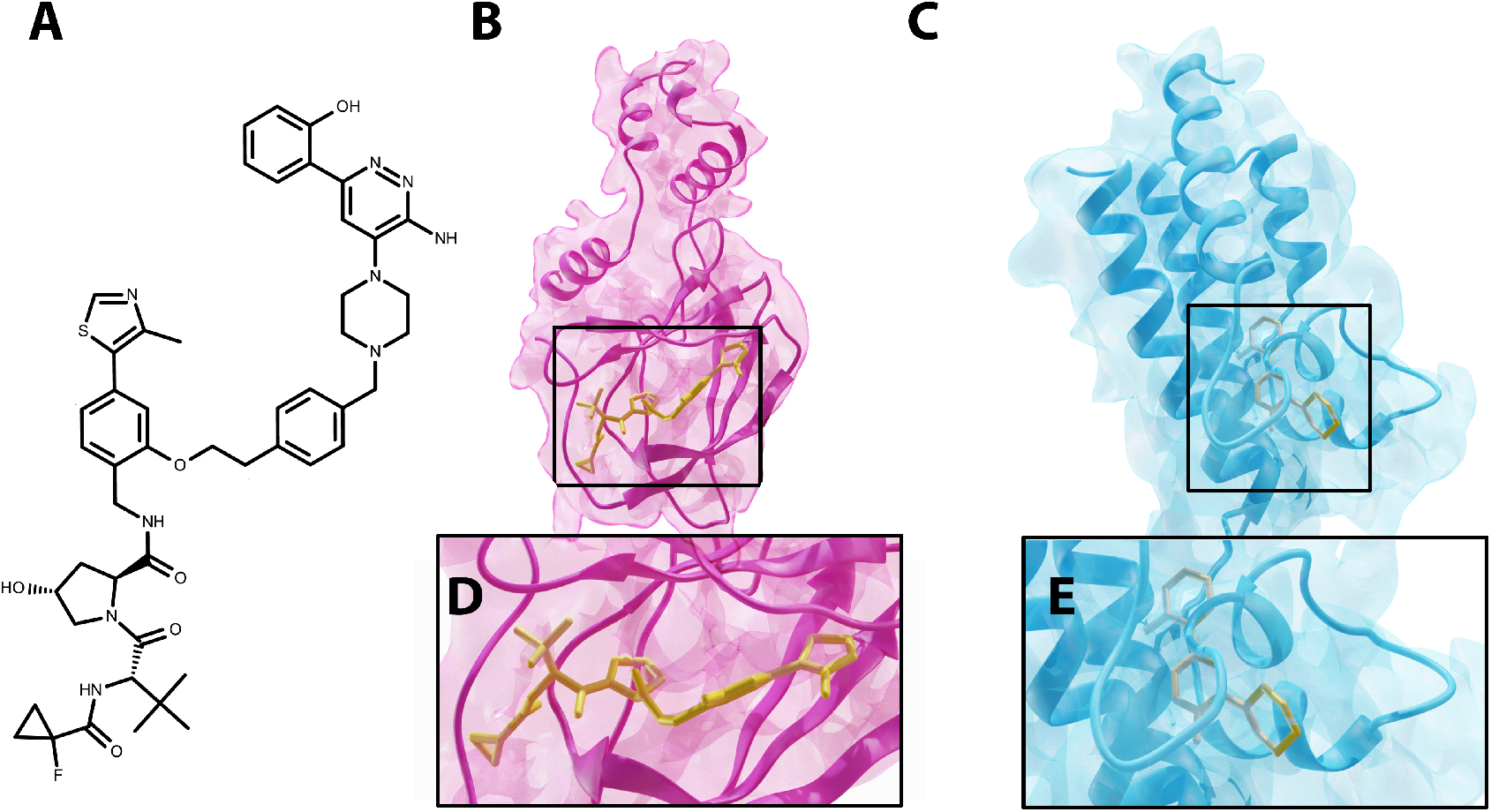
Example of the required inputs for the ternary complex 6HAX (PDB ID). (A) PROTAC molecule. (B) and (D) E3 ligase docked to warhead. (C) and (E) POI docked to the warhead.

It is possible to apply our method to the structures of proteins and docked ligands extracted from the co-crystal structure of known PROTAC ternary complexes (so-called bound protein data). However, as we do not have access to these bound structures in most real-world situations, we focus on using unbound data (individual protein structures). Note that this makes the problem more challenging as the 3D structures in the bound and unbound cases differ.

### 2.2 Bayesian optimization

Our goal is to optimize a black box function *y*(**x**) describing the quality of a ternary complex pose (defined by the vector **x**), where evaluations are expensive and result in noisy observations. In the present work, the noisy observations result from an objective function that combines information concerning the favorability of protein-protein interaction with information on the feasibility of finding a stable PROTAC conformer for a particular ternary complex **x**. This is a classical use case for Bayesian optimization [45, 46]. Formally, our goal is to find the optimal pose **x**\* which maximizes *y*(**x**), i.e.,

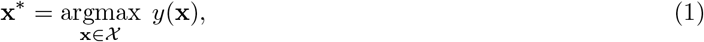

while searching within some conformation sample space ***χ***. This is accomplished by using the noisy observations obtained by evaluating the objective to update a surrogate model (here, a Gaussian process), which is used to sample promising points for evaluation using an acquisition strategy. These steps are repeated in a loop (Figure 3) until some stopping criterion is met. The loop is described in algorithm 1. In the following sections, we describe the elements of this loop (search space, objective evaluation, surrogate model, acquisition strategy) in detail.

**Figure 3:**
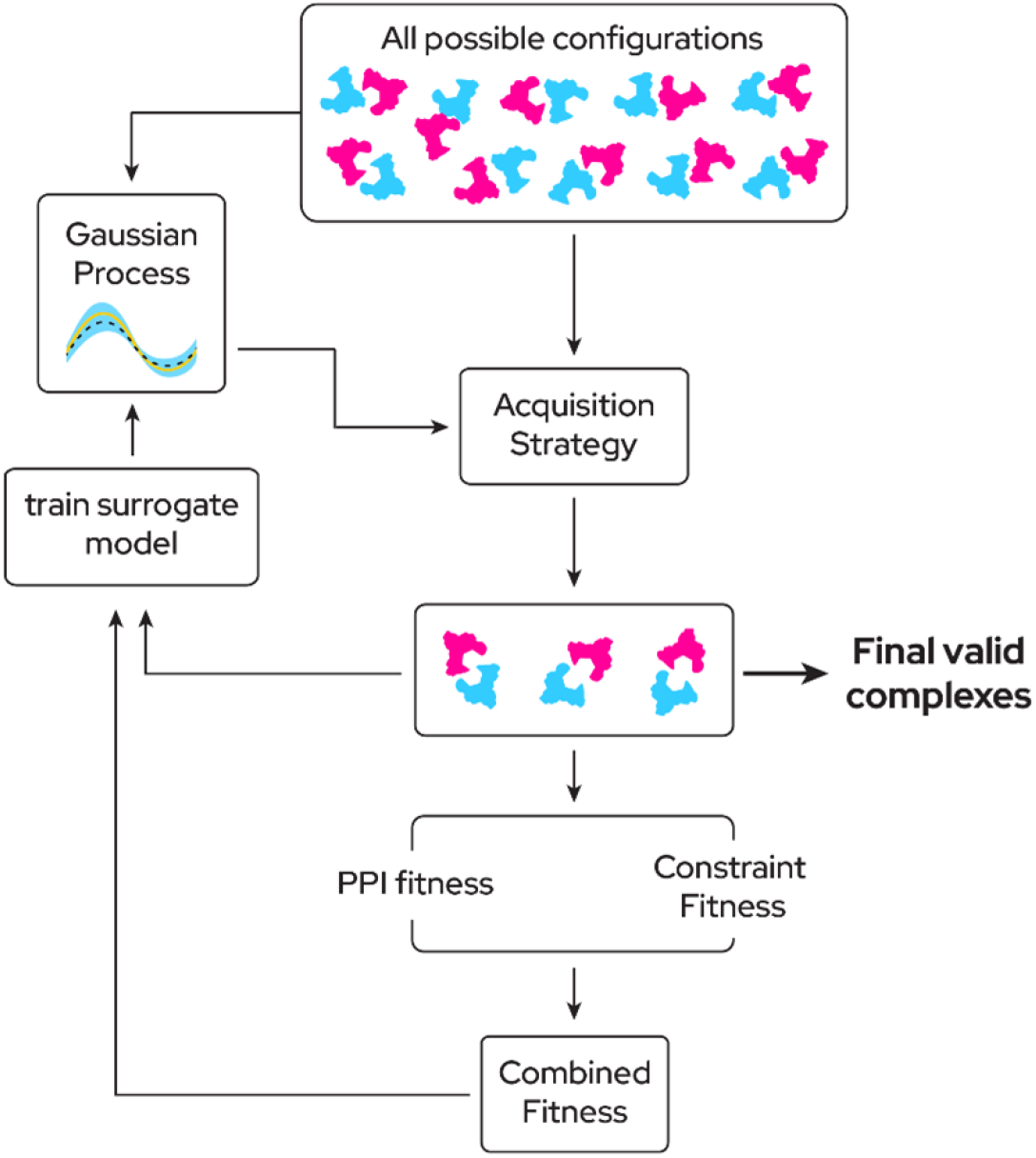
Overview of Bayesian optimization pipeline. We are sampling from the roto-translational space of POI. Then we assign a score for each sample *D* = {(**x**_*i*_,*y_i_*)}. Then, the dataset *D* is used for updating the GP into posterior GP. For sampling the next batch of points, the acquisition function is used. In our proposed method, we used the UCB acquisition function to determine the next sampling points. This iterative process continues until the end of some Bayesian optimization steps. The number of iterations is a hyperparameter that controls when do we get the list of final valid complexes.

#### 2.2.1 Search space

Using the Bayesian optimization loop, we optimize the ternary complex pose. Each such pose is described by a vector **x**. As we are concerned with the relative orientation of the two proteins, we arbitrarily fix the E3 ligase at the origin and only modify the position of the POI while attempting to find the pose with the highest fitness. The vector **x** thus describes the relative rotation and translation (RRT) of the POI with respect to the E3 ligase, and comes from the 7D roto-translational space ***χ*** (3 dimensions for translation, 4 dimensions for rotation as a quaternion). In the remainder of this paper, we refer to such protein-protein configurations as RRTs (see Figure 4A for an illustration). Note that the alternative of optimizing using an all-atom representations for the POI would also be possible (e.g., encoding the location of each residue within **x** to allow the optimization algorithm to also alter the protein structure), but the resulting search space is less suitable for practical Bayesian optimization due to its high dimensionality.

**Figure 4:**
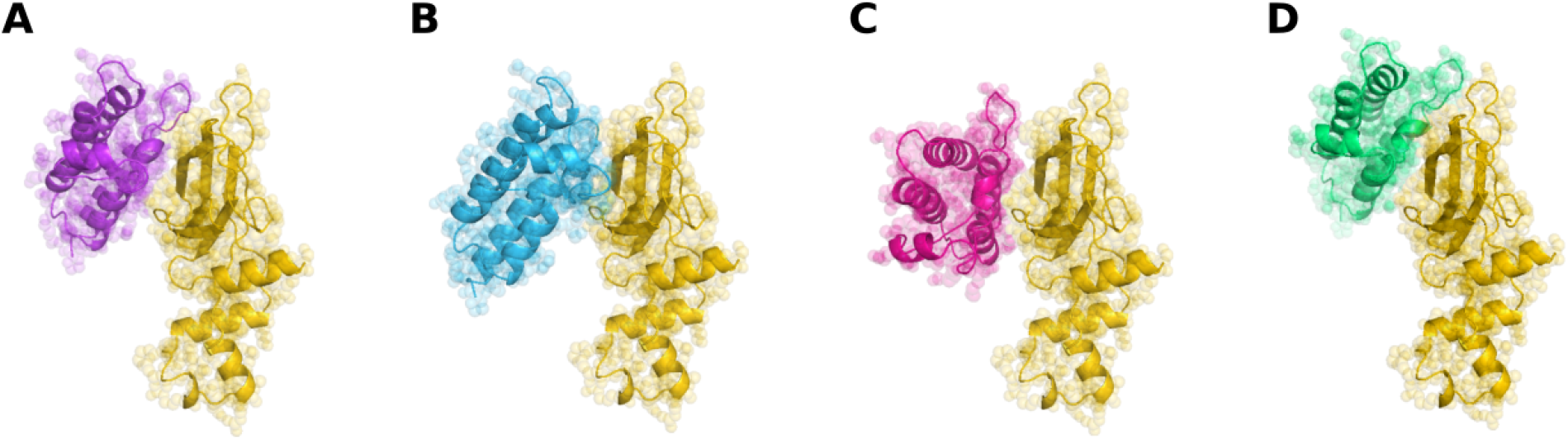
Illustration of the search space. The E3 ligase (yellow) is fixed in space. The Bayesian optimization optimizes the ternary complex by sampling different poses (A-D), each of which describes the rotation and relative translation (RRT) of the POI.

#### 2.2.2 Objective evaluation

Our model uses Bayesian optimization to propose promising candidates **x** ∈ ***χ*** for evaluation. This requires training a surrogate model using the currently available data, i.e., a dataset *D* = {(**x**_*i*_,*y_i_*) | *i* = 1, 2, &,*N*} consisting of RRTs **x**_*i*_ with their assigned fitness values *y_i_*. We used a combined with two components:

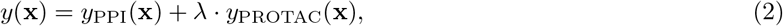

where λ determines the relative weighting of the PPI fitness *y*_PPI_ vs. the PROTAC fitness *y*_PROTAC_ (we used λ = 1). The PPI fitness represents the strength of interaction between the two proteins. While strong PPIs are important for a stable ternary complex, there are cases in which the proteins are oriented such that the respective binding pockets for the E3-binder and the warhead are too far apart for the length of the linker. On the other hand, the proteins’ relative positions might result in binder and warhead being too close, leading to steric clashes within the linker itself. The PROTAC fitness *y*_PROTAC_(**x**) ensures these cases have a low overall fitness values.

**Algorithm 1.**
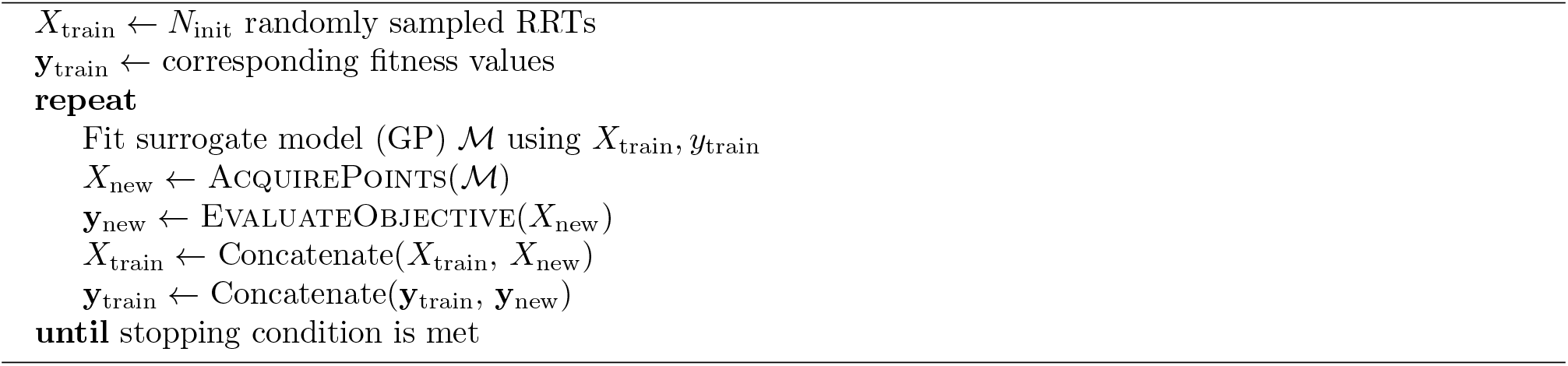
The BO loop

**Algorithm 2.**
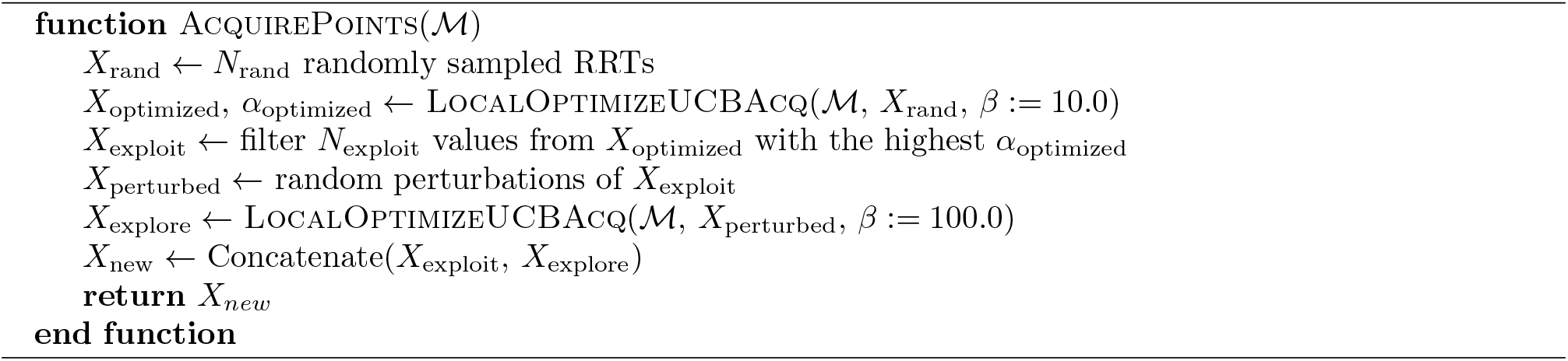
Acquiring new points for evaluation

**Algorithm 3.**
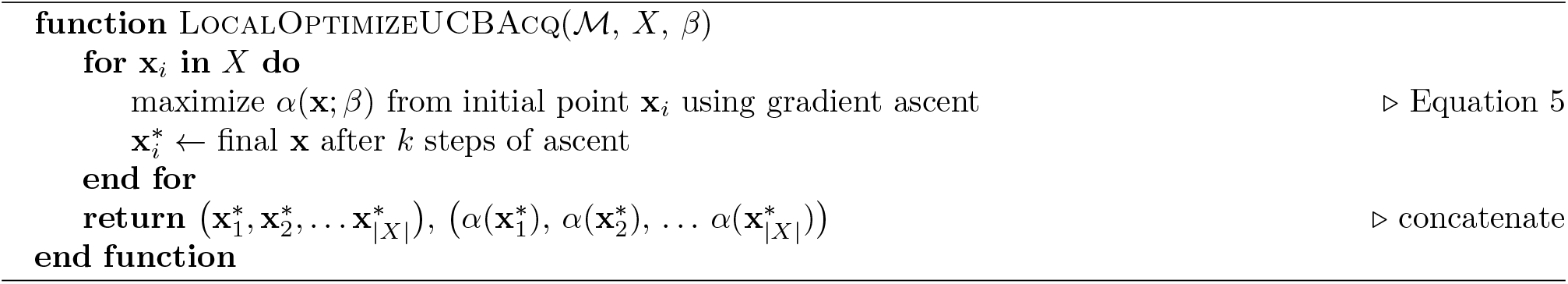
Acquisition optimization algorithms

**Algorithm 4.**
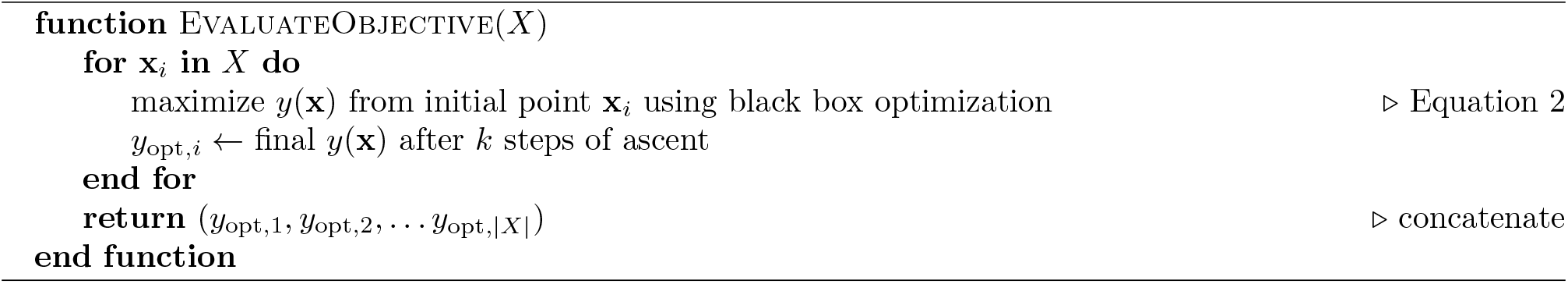
Evaluation of objective

##### PPI fitness

A number of protein-protein docking scoring schemes have been proposed in the literature. In this work, we use the distance-scale finite ideal-gas reference (DFIRE) energy score [47] as *y*_PPI_. This score is a relatively unbiased estimate of the validity of the protein-protein interaction while allowing rapid evaluation. The PPI fitness serves to filter out infeasible protein-protein orientations.

##### PROTAC fitness

The PROTAC fitness *y*_PROTAC_ is calculated based on a linker conformer, which we generate using RDKit. To do this, we constrain the two binding fragments are at their respective target positions (specified by the protein orientations as described by **x**) via a restoring force field. The linker atoms are free from any such constraint, and we sample linker conformations by minimizing the UFF [48] starting with initial conformers generated using distance geometry. After generating 4 candidate conformers, we choose the output to be the conformer with the lowest RMSD between the fragments’ positions and their target positions. The PROTAC fitness is defined as the negative square-root of the UFF energy of this conformer.

The generation of this conformer and hence fitness is computationally expensive. We found that we can effectively train a neural network model to effectively predict the PROTAC fitness. For a particular POI, E3 ligase, and PROTAC, we train a fitness model which maps the 7D RRT vector x to the PROTAC fitness values generated using RDKit (as described above). The model consisted of two components: a simple model which learns to predict asymptotic component of the energy (based on the distance between the translation component of the RRT and a learned center vector), and a 3-hidden layer neural network which learns to predict the residual component. Prior to the BO loop, we pre-train this model using a dataset of RDKit-generated conformers and use it to map the x values proposed by the Bayesian optimization loop. The advantage is that after learning the model, for computing the fitness for new rotation and translation, just one forward pass is required, making the fitness computation very fast.

It is important to note that the above PROTAC fitness values only use the protein geometry in order to define the fragment positions. For the linker atom positions, the proteins are not taken into account, thus, it cannot be used to filter out cases with steric clashes between PROTAC and proteins. These cases are taken into account by the PROTAC stability score (see below).

##### Local fitness ascent

When evaluating the objective, we additionally perform a local ascent in the space of RRTs using a black box optimization algorithm (Nelder-mead simplex ascent [49]) and take the final optimized objective as the objective corresponding to a particular orientation x. This gives a more robust estimation of the objective as in some cases, small changes of x can have a somewhat large influence on the fitness. This approach is described in algorithm 4.

#### 2.2.3 Surrogate model

We aim to optimize the black box function *y* (**x**) using the evaluations *y_i_* at some points **x**_*i*_ but without having access to an analytical form of this function. To optimize y, we need to approximate/extrapolate our belief about y at points that have not been evaluated in order to find regions of interest within the search space ***χ***. This is done using a surrogate model *f* (**x**), which describes our belief/approximation of the actual black box function *y*(**x**) based on our observed points so far. The surrogate model is trained so that it matches the true function y at the evaluated points **x**_*i*_, but it can also be evaluated at other points at a low computational cost. The important characteristic of the surrogate model is the ability to output uncertainty of the prediction in the form of posterior over function values *f* (**x**) at points **x**.

While in principle Bayesian optimization can utilize different surrogate models, we employed the widely used Gaussian process (GP) [50, 51]. A GP is defined by a prior mean function m(**x**) and a covariance kernel *k*(**x, x**’) which describes the covariance between any two points **x** and **x**’, i.e.,

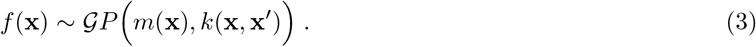

We use a constant learned value *m*(**x**) ≡ *m*_0_ as prior mean. As covariance kernel, we use the a well-known Matern kernel [51] (see Figure 5), i.e.,

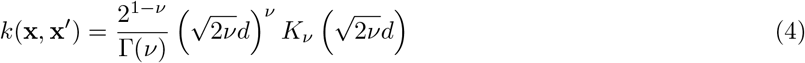

where *d* is the distance between **x** and **x**’, *v* is a smoothness parameter (taking on values 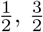, or 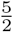, with larger values increasing smoothness), and *K_ν_* is a modified Bessel function. As distance *d*, we use the RMSD between the atomic coordinates of the protein of interest (scaled by a learned parameter parameter Θ), which (between two rigid transformations of an object) can be calculated very efficiently, making this kernel computationally inexpensive.

**Figure 5:**
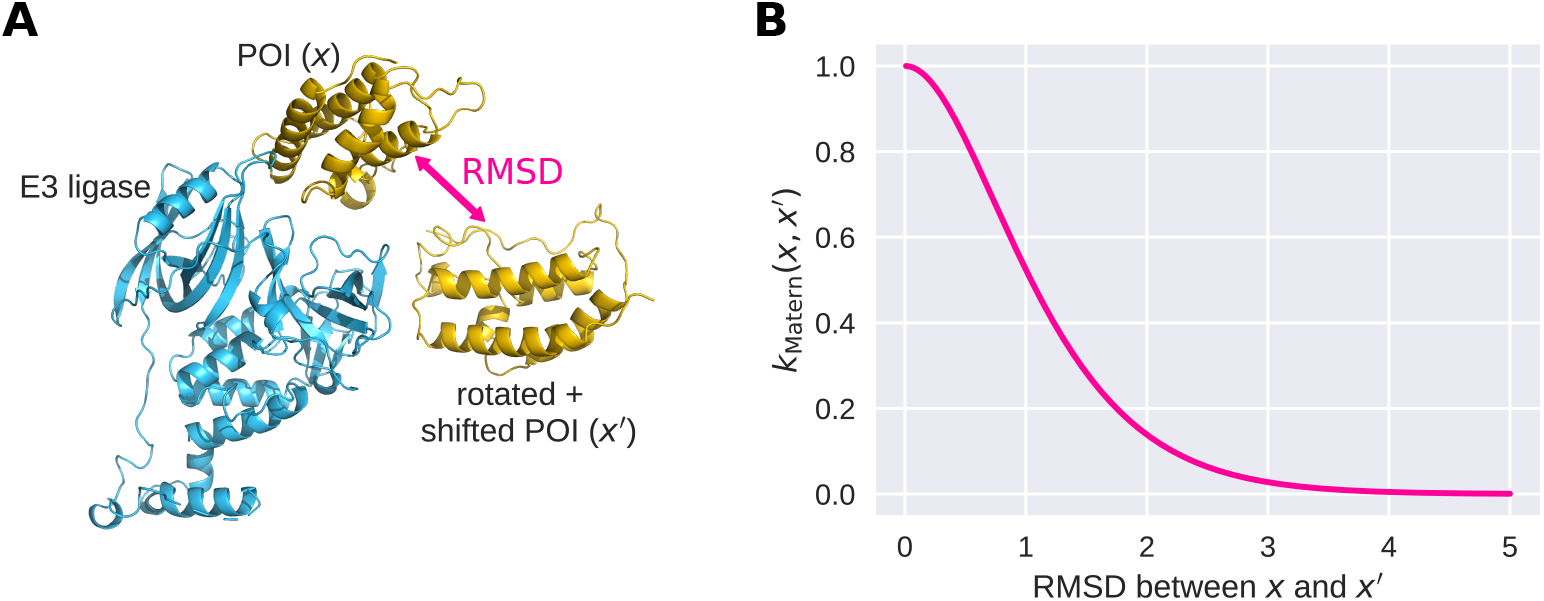
Illustration of the Matern kernel, used as covariance kernel for the Gaussian process. (A) RMSD between different POI positions relative to the E3 ligase (i.e., different RRTs **x**). (B) Matern kernel (see Equation 4) as a function of the RMSD.

#### 2.2.4 Acquisition strategy

Bayesian optimization requires a strategy by which we select new candidate RRTs for which we evaluate the objective. Intuitively, we want to perform exploration by sampling points for which we have no information about their potential value, while also performing exploitation by sampling points which we assume will have a high objective value. At the same time, we should avoid sampling points where we already possess enough information to predict that the objective will be low. Formally, we define an acquisition function *α*, and evaluate the objective on values **x** which maximize *α*. There are different acquisition strategies allowing to balance exploration and exploitation [35]. We found that the popular upper confidence bound (UCB) acquisition function [52, 35] is effective in our scenario. It is defined as

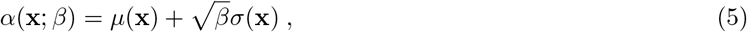

where *μ*(**x**) and *σ*(**x**) are the predicted mean and standard deviation of *f* (**x**) (which can be calculated from the existing data points using the mean function and covariance kernel). The parameter *β* > 0 defines the balance of exploration versus exploitation (a large *β* leads to large *α* at points where the standard deviation is high, thus increasing the exploration, while a small *β* leads to large *α* at points with a high predicted mean, thus increasing exploitation).

Our full acquisition strategy is described in algorithms 2 and 3. In each iteration of the BO loop, we sample two batches of new points, one for exploitation and for exploration. We do this by selecting points with a high value of the above acquisition function, with low and high *β* values respectively, and performing local optimization using gradient ascent (using ADAM [53]). We use the open-source BoTorch package [54] to solve this optimization problem and implement the BO loop.

### 2.3 Local optimization with simulated annealing of the PROTAC stability score

From the above points, we filter out RRTs with infeasible PROTAC scores or PPI scores. We next locally optimized the remaining RRTs using a combination of the PPI score and a PROTAC stability score.

#### PROTAC stability score

We calculated a PROTAC stability score using an extension of AutoDock-Vina [55]. This score takes into account the stability of the PROTAC within the context of the protein configuration corresponding to a particular RRT (which is ignored in the calculation of the PROTAC fitness, see above). We extended Autodock-Vina by adding an additional restoring force that allows constraining some atoms of the ligand while performing docking. We used this force to keep the E3 binder and the warhead at their respective coordinates given the current protein-protein RRT, ensuring that the binding fragments do not leave their binding pockets. We calculated ad PROTAC stability score for each sampled RRT by performing the following steps:

1. We optimized the PROTAC molecule’s conformation under only the restoring force and the intra-molecular part of the Vina force field, creating a viable conformer with the binding fragments held at their positions.
2. We then turned off the restoring force and perform an optimization using the full (intra- and inter- molecular) Vina force field.

For each RRT, Vina creates 10 (random) PROTAC conformers and performs this optimization (using the sum of the energies of the restoring force and the Vina force fields as fitness). Correspondingly, for each such PROTAC conformer *i* we have the following two energies. First, the Vina energy 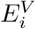, which is the combination of the intermolecular and internal energy of the conformer. Second, the restoring energy, 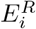 which measures the deviation of the E3 binder and warhead from their constrained positions in the particular RRT. To generate a score corresponding to the stability of the PROTAC in a particular protein-protein RRT, we look at the following cases:

1. 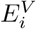 is low and 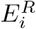 is low: the conformation is stable and doesn’t strain the warhead and E3 ligase.
2. 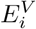 is low and 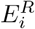 is high: the conformation is stable, but the warhead and E3 ligase need to be displaced to achieve this conformation.
3. 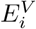 is high: the conformation is unstable/infeasible.

We thus define the following weighted average of the Vina energy as the PROTAC stability score:

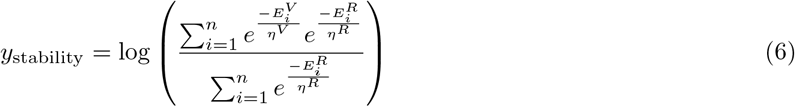

where *η^v^* and *η^R^* are scaling parameters. This score rewards stable conformers while being forgiving to outliers.

#### Simulated annealing

We used a combination of this score and the PPI score to perform local optimization of the remaining candidates using simulated annealing [56]. For any RRT, simulated annealing proceeds via a sequence of steps, where in each step we randomly sample an RRT near the current RRT. The samples generated here are optimized to have a reasonable PPI and PROTAC score. We evaluate the above mentioned weighted sum of the PROTAC stability and PPI scores and stochastically accept and reject it based on that.

As this process allows the PROTAC conformation to change (both explicitly through simulated annealing and implicitly within Autodock-Vina), it resembles a refinement of the PROTAC conformer.

Finally, we ensure that each sampled point has a reasonable PPI and PROTAC score.

### 2.4 Clustering and filtering using TCP-AIR energy

After generating this refined set of candidate RRTs, we perform a clustering step. These clusters are first filtered using the TCP-AIR energy, then, we re-rank them to obtain the final output set of RRTs.

#### Clustering

We cluster the set of candidates using a combination of the PPI score and the PROTAC stability score. Clusters were greedily created by selecting the RRT with the highest fitness, choosing that as the center of the new cluster, and assigning all sampled RRTs with an RMSD (vs. the center of the cluster) of below 5 Å to this cluster. This step was repeated (starting again with the RRT with the highest fitness which was not yet part of any cluster) until all points were assigned to some cluster. The motivation for this simple clustering strategy is that it guarantees a limit on the maximum RMSD between the poses assigned to a single cluster. It is to be noted here that clusters with a single element are also considered.

#### Filtering using TCP-AIR energy

We use a filtering technique based on the ambiguous interaction restraints (AIRs) energy [57] that has been used extensively in [36] for energy calculation. In order to calculate the TCP-AIR energy terms, for both proteins, active and passive residues need to be defined. Active and passive residues are chosen based on two criteria: first, they have some atoms on the surface of the protein, and second, these atoms lie within some distance of the atoms of the fragments binding to this particular protein. We used distances of 5 Å and 7.5 Å for active and passive distances, respectively. For instance, residues within 5 Å of the warhead were defined as active residues for POI. For these residues, we calculate the effective distances (as done in previous work [36]). These are used to calculate the TCP-AIR energy (using the NOE energy formula, [57]) using three threshold values *L, U*, and *S. L* and *U* are the lower and upper bounds on the effective distance, while *S* is the value from where the computation of the energy changes from quadratic to linear. These parameters need to be specified for each ternary complex separately since each structure has different distances between two active sites of the proteins (due to the space occupied by the PROTAC molecule). We set the values, we calculated the average distances of each active residue of one protein with the active and passive residues of the other protein, resulting in two distances for each candidate. We considered all of our samples that have PROTAC stability score > 5. For each of them we took an average of the two distances of the proteins and define L as the 25th quantile. We then defined *U* = *L* + 3 Å and *S* = *U* + 2 Å. When filtering, we preserved all clusters that had at least one candidate with an TCP-AIR energy large than the 90th quantile of all AIR energies (see Figure 6).

**Figure 6:**
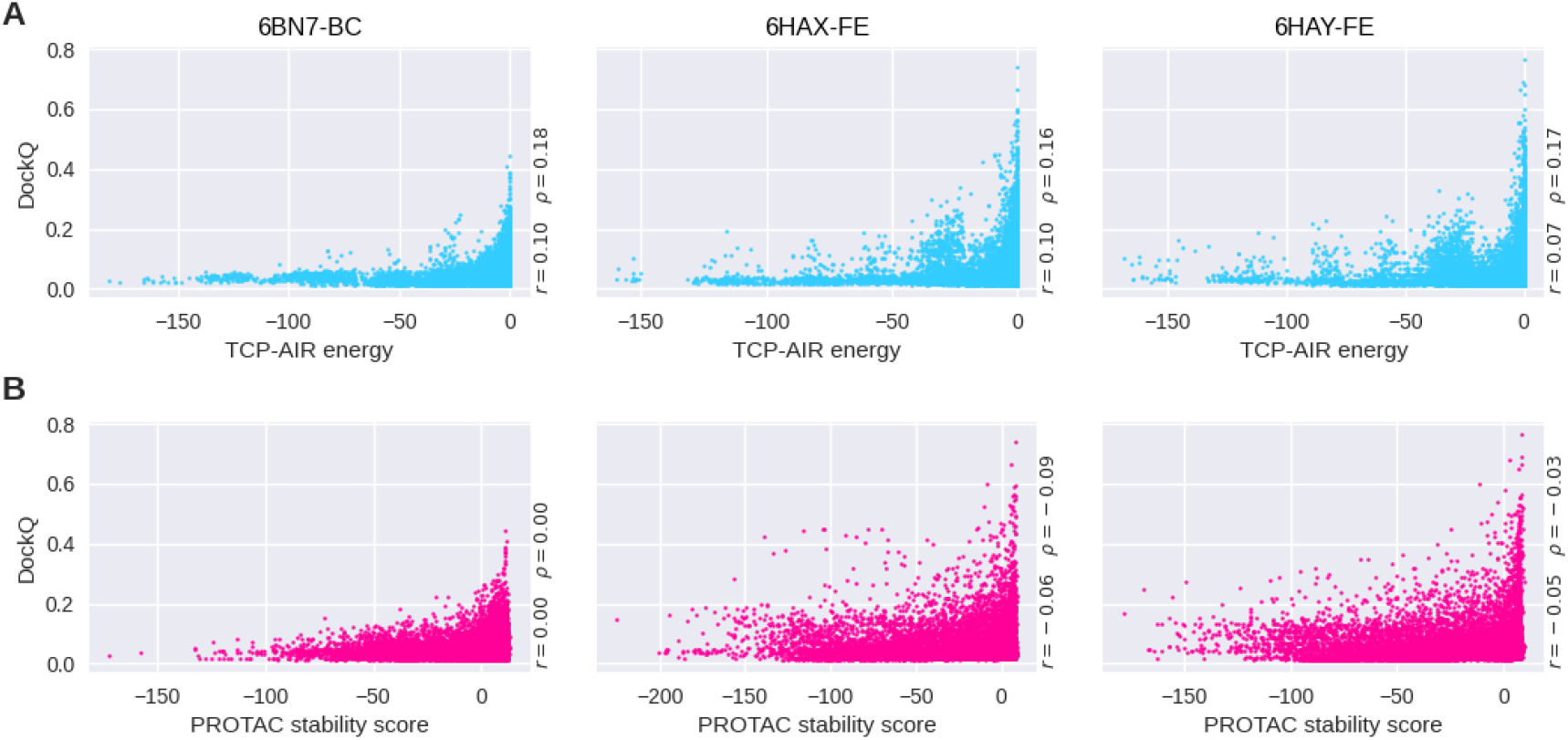
Usefulness of TCP-AIR energy and the PROTAC stability score for filtering. A) Correlation between the TCP-AIR energy and the DockQ score for different example ternary complexes. As high DockQ scores are only achieved when the TCP-AIR energy is 0 or close to 0, the later can be used as a filtering criterion. Text insets show Pearson (*r*) and Spearman (*ρ*) correlation values. B) Same as A, but for the PROTAC stability score.

#### Re-rank clustering

After filtering, we re-rank the remaining clusters using the PROTAC stability score (see above, Section 2.3). Clusters are ranked according to the largest PROTAC stability score of their members. We then calculate the rank of the near-native clusters (Table 2). For the refinement, we generate a diverse set of complexes by selecting the structure with the highest PROTAC stability score out of each of the 100 top-ranked clusters. For each of these, we use the 10 PROTAC conformers generated by AutoDock-Vina (see Section 2.3) as input to the refinement (i.e., the refinement is performed for 1000 structures).

### 2.5 Refinement

Starting with a given number of ternary complex structures from re-ranked and clustered TCP output, we performed further refinement using classical force field-based molecular simulation with an implicit solvent model. Force field parameters for each molecule were taken from the Amber ffSB14 force field [58] for proteins, and GAFF2 [59] for PROTAC molecules. Partial charges for the PROTAC molecules were determined as AM1BCC charges [60] using antechamber/Ambertools version 21 [61]. MD simulations were performed using OpenMM [62] with a Langevin type integrator and a time step of two femtoseconds. The implicit solvent model used was the second GBneck model (GBN2 [63]). Each structure was first optimized through energy minimization for 100 steps to remove close contacts, followed by a simulated annealing run, with the temperature varying from 300 to 0 Kelvin in a linear fashion, for a total simulation time of 200 picoseconds. This procedure was repeated four times to approximate the average total free energy of solvation and standard deviations for each input structure. The resulting energies are essentially a simplified version of MM-GB/SA energies, where a calculation of entropic contributions is omitted. Including entropy terms would require much longer simulation times, and, given the approximate nature of current approaches for entropy estimation [64], we consider the benefit of including this contribution limited.

### 2.6 Re-clustering and Re-ranking

The refined structures (1000 of them) were clustered by the fraction of common contact (FCC) clustering method [65]. Afterwards, the clusters were ranked based on size (the cluster with the highest number of elements being the first-ranked). The clustering threshold was set to 0.9 (this could lead to smaller entropy in the clusters [65]) and minimum number of elements in a cluster was set to 4. The rest of non-clustered structures were not taken into consideration.

### 2.7 Fitness landscape analysis

Using 5 exemplary ternary complexes from PDB (bound structures), we extracted the RRT corresponding to the native pose and modified them to investigate the resulting change of the fitness *y*(**x**). For each ternary complex, we generated 4000 random near-native samples by first randomly shifting the POI before rotating them. The shift directions were drawn from a von Mises-Fisher distribution with the mean set to the offset of the POI (i.e., pointing away from the E3 ligase) and the directional parameter *κ* = 1 (i.e., shifts moderately concentrated in this direction). The shift distances were drawn from an exponential distribution with λ = 1 Å. The rotations were generated by drawing a random rotation direction from a uniform distribution (no directions favored) and a rotation angle drawn from a von Mises distribution with mean 0 and *κ* =10 (i.e., small rotations favored).

## 3 Results

### 3.1 Evaluation of fitness functions

To check whether our proposed fitness functions are a meaningful objective for the BO loop, we analyzed the change of the fitness values around the true ternary complexes. Using bound structures, we created random near-native RRTs by altering the position of the POI using small, random translations and rotations (see Section 2.7 for details). We then compared the values of the PPI and PROTAC fitnesses of these random RRTs to the fitness of the actual native complex (Figure 7). The results show that these fitnesses are higher at the native RRT compared to other RRTs in its vicinity, thus indicating the their usefulness as objective for the BO loop.

**Figure 7:**
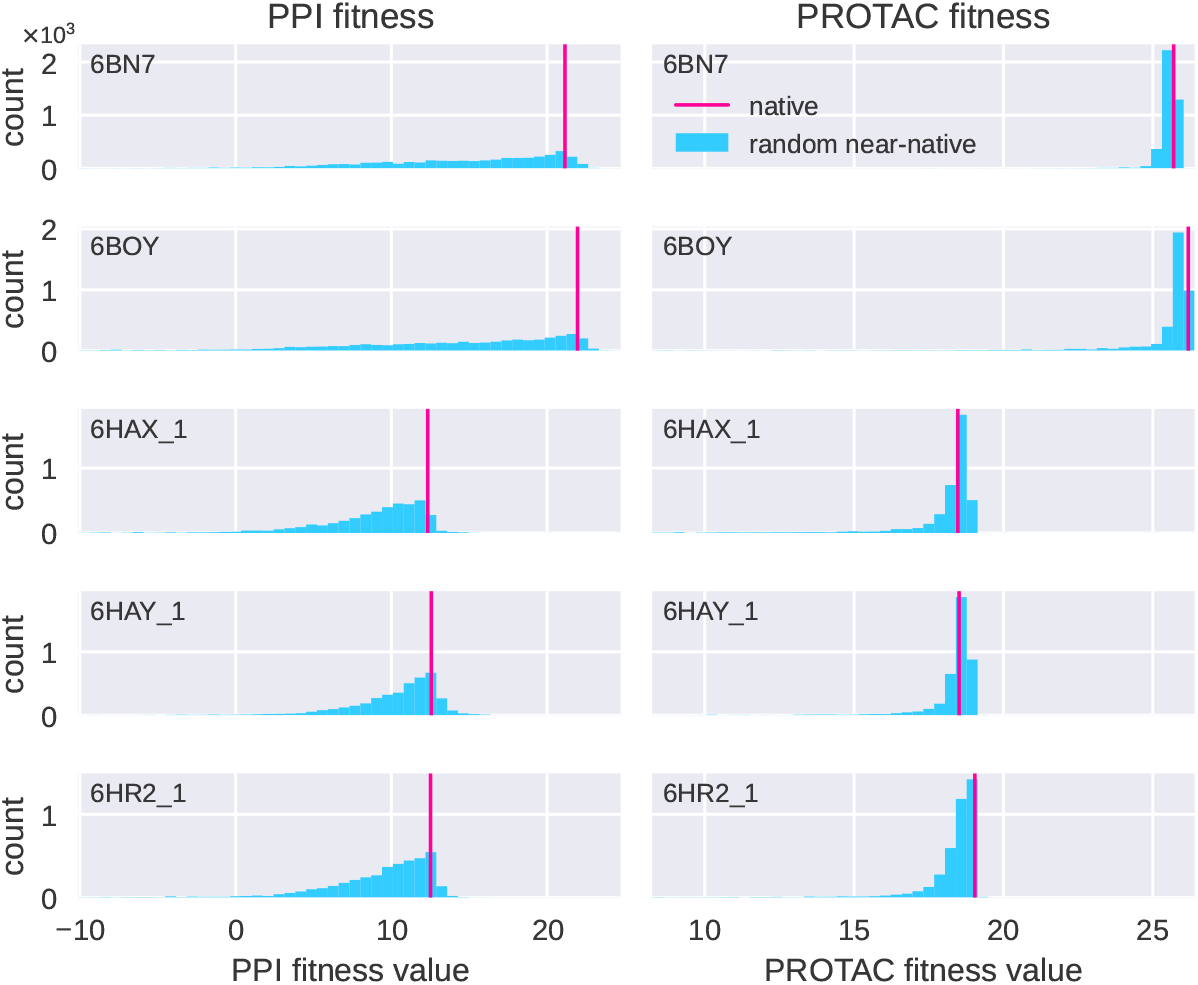
Native complexes have high fitness values. The columns show (for five sample ternary complexes) histograms of PPI fitness (left) and PROTAC fitness (right) for random near-native RRTs, created by randomly shifting and rotationing the POI (see text for details). The actual native complex (pink line) achieves a comparably high fitness.

### 3.2 Evaluation of the BO loop

We futhermore evaluated the BO method to test whether it performs adequate exploration and exploitation. Ideally, the distribution of candidates acquired by the BO loop is clustered in regions where the fitness *y*(**x**) is high. We compared the BO outputs to a set of randomly sampled candidates, and found that this is indeed the case (Figure 8): while random sampling occasionally finds RRTs with a high fitness, the BO loop generates samples in a more structured way (Figure 8A-B), resulting in more consistently high fitnesses (Figure 8) and a higher maximum value.

**Figure 8:**
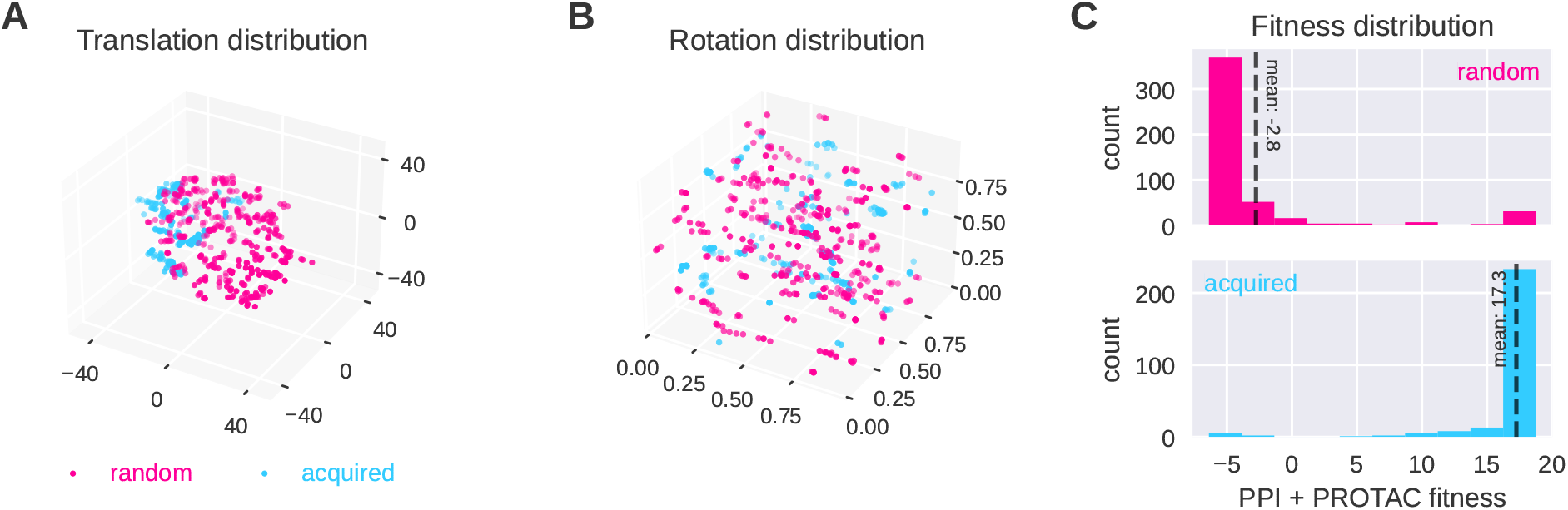
Bayesian optimization results in higher fitness values than random search. Shown here are samples from the acquisition function at iteration 15 of the BO loop for the 6HAX complex, compared to random samples picked with proteins close to each other. (A) Translation values from random sampling (pink) and proposed by Bayesian optimization (blue). The latter leads to a candidate concentration in a promising region. The spread of points also suggests that the the BO loop sufficiently explores the space without converging prematurely to any specific minima. Points sampled via Both random sampling and BO are subject to a local optimization (B) Rotation quaternions mapped onto a 3-D cube from random sampling and Bayesian optimization. (C) Resulting fitness histograms. While random samples find some conformations with high fitness, the best fitness acquired by the Bayesian optimization is higher, and the mean fitness is substantially better, confirming that our samples efficiently search through the space of higher fitness values.

### 3.3 Evaluation metric

For evaluating the quality of the candidates generated by the BO loop (Table 2), we used the DockQ score [66], which is based on the metrics underlying the well-known critical assessment of predicted interactions (CAPRI) score [67]: *f*_nat_, *I*_RMSD_, and *L*_RMSD_. *f*_nat_ is the fraction of native contacts that a candidate recovers. *I*_RMSD_ is calculated over all backbone atoms for those residues found in the interface of the reference structure (residues within a radius of 5 Å). *L*_RMSD_ is calculated based on the backbone atoms of the ligand between the predicted and native model (after aligning the receptor structures, i.e., the E3 ligase). The DockQ score combines these measures into a single scalar value between 0 and 1, with higher values corresponding to ternary complexes which resemble the original structure more closely. The distribution of the DockQ scores has been defined as follows [66]:

- 0 < DockQ < 0.23 - Incorrect complex
- 0.23 ≤ DockQ < 0.49 - Acceptable quality complex
- 0.49 ≤ DockQ < 0.80 - Medium quality complex
- DockQ ≥ 0.80 - High quality complex

### 3.4 Prediction of near-native poses

We run our pipeline on 14 ternary complexes, some of which have multiple reported native ternary complex structures leading to a total of 22 cases. By running our pipeline and comparing our results to these 22 native ternary complex structures, we have found that the BO loop can effectively sample the near-native poses. In addition, the filtering and ranking procedures can assign them a relevant rank. For 21 out of 22 cases, we could sample at least acceptable quality complexes. For 16 out of 22 (Table 2), we were able to rank them within the top 20 clusters, and for 12 out of 22 complexes, we were able to recover a medium quality ternary structure. Additionally, 14 out of 22 clusters have at least 50% of the near-native poses in the reported cluster (see Table 2). This method proves that even with using a relatively simple PPI interaction and unbound structures, taking into account the constraints imposed by the linker, with the incorporation of the PROTAC score in the BO loop, as well as the TCP-AIR energy for filtering and PROTAC stability score for ranking, we can recover near-native predictions of the ternary complex structures.

After the use of refinement, we can see clear improvements. For 22/22 cases, we obtained complexes with the DockQ ≥ 0.23. For 17 out of 22 (Table 3), we were able to rank them within the top 20 clusters. For 14 out of 22 cases, we could recover at least a medium-quality ternary structure. For 2 out of 22 complexes, we recovered a high-quality ternary structure. Additionally, in 14/22 cases, in the reported highly-ranked near-native cluster, at least 50% of the poses in it are near-native poses (see Table 3).

**Table 3:**
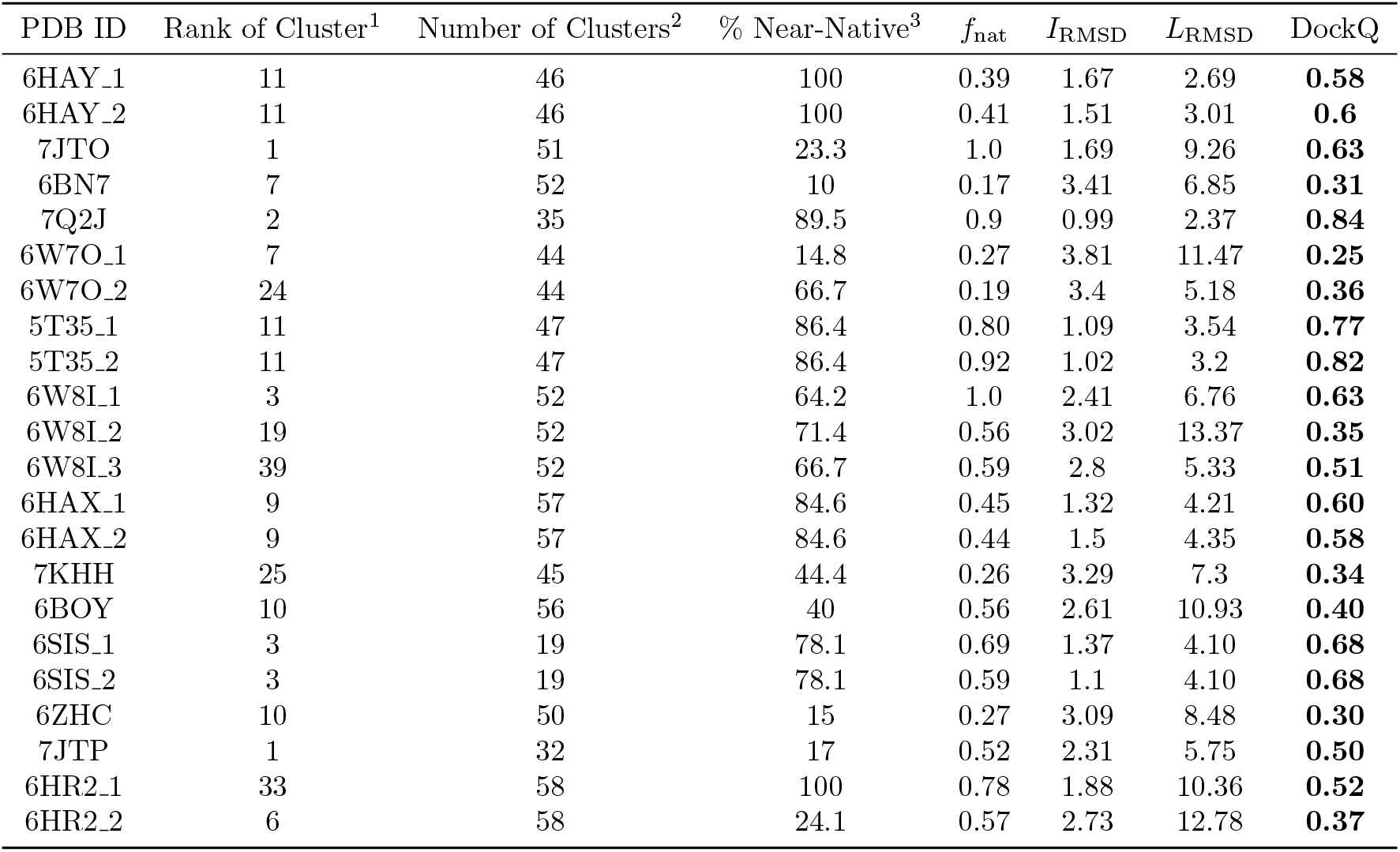
Results of the BOTCP method for ternary complex prediction on unbound structures after the refinement. ^1^The best rank containing at least one model with DockQ≥0.23. ^2^The total number of clusters. ^3^The percentage of the near-native poses in that specific ranked cluster. For each ternary complex, we selected a cluster which includes an RRT with a high DockQ score. Columns also show the PDB ID and characterization of the best RRT in terms of *f*_nat_, *I*_RMSD_, *L*_RMSD_ and DockQ score (see text).

### 3.5 Comparison to previous work

Ternary complex prediction models have frequently been evaluated on bound structures (e.g., PRosettaC [32]). We have demonstrated in this work that effective ranking of predicted structures in the more challenging scenario of unbound structures is possible with our method. We are aware of one other approach based on unbound structures has been presented by Weng et al. [33], who have shown successful ranking of the near-native clusters within the top-15 clusters. However, Weng et al. used clustering with FCC using a threshold of 0.5, which is relatively low, leading to a large spread of poses within the resulting clusters. Even though there may be a near-native pose in such a cluster, a threshold of 0.5 implies that the fraction of near-native poses in the cluster would be low due to the diversity of poses within the cluster. Moreover, there is a higher likelihood that the best-scored pose in this cluster is, in fact, not a near-native pose. To avoid this, we choose a higher threshold of 0.9, which restricts the spread of points placed in a single cluster. Accordingly, a cluster ranked high in our case has lower entropy, i.e., the yield of near-native structures in it is much higher. With this form of constrained clustering, we have near-native clusters ranked within the top-15 clusters in 13 out of 22 cases before refinement and 16 out of 22 cases after refinement. These results demonstrate the efficacy of the Vina and TCP-AIR energies.

## 4 Conclusion and outlook

We have demonstrated our BOTCP module for predicting unbound rigid ternary complex structures (Table 2). Our method can sample and appropriately rank the near-native poses for most of them. Initially, 7KHH was the only complex for which our method is (currently) unable to sample near-native poses effectively. However, after refinement, we were able to recover near-native poses for all of the complexes (Table 3). Our results do not require the use of GPUs to compute the scores and, using 128 standard x64 CPU cores, for each complex, our results including filtering and re-ranking excluding refinement, typically take less than 2 hours. This represents a significant improvement over conventional techniques, that attempt to handle many interactions in a ternary complex via GPU intensive molecular dynamics simulations which typically take days or weeks to report a near-native pose. Note that this code is a prototype with significant room to optimize the various components further still.

At the heart of this, is the fact that Bayesian optimization learns to sample regions of high scores in a few iterations. In all of our simulations, we sample only around 2500 points, where the BO loop converges rougly after sampling around 700 points. This sample efficiency suggests modifying the fitness function to use more computationally expensive but informative data (e.g., the PROTAC stability score) for the BO loop. Such a fitness could take into account the interactions of the linker with the proteins, which has the potential to significantly improve the yield of near-native poses.

Currently, there are several avenues of improvement for both the quality of results and computational efficiency. The subsequent refinement of the structures provided from the BOTCP module currently takes about 25 hours but could be further optimized. For instance by running the refinement for less than 1000 poses or by using a smaller number of PROTAC conformations per structure. While the refinement is still computationally intensive, we see that the results without refinement suggest that the current scores and sampling technique can be beneficial in assessing the interactions for a given ternary complex.

As mentioned above, incorporating more informative PPI and constraint scores in the BO loop will further filter non-native poses. Currently, the PROTAC stability score and the PPI score are affected because the side-chains are misaligned with the side chains as in the original crystal structure. Our experiments show that the native structure consistently has a PROTAC stability fitness that is among the highest observed for that complex, which promises significant improvements if this score includes side-chain flexibility (e.g., using Autodock-Vina). To summarize, we demonstrate in this work the successful application of Bayesian optimization as well as the design of two specialized scores that can effectively sample and rank highly clusters of near-native poses for Ternary Complexes.

## Author Contributions

A.R., T.T. conceived the algorithms, and ran the simulations using the TCP-AIR score, PROTAC stability score and the BO pipeline. A.R., T.T., M.M., H.F. implemented the software necessary. M.B. performed input data processing and refinement of structures. M.M. performed analysis of the fitness scores and result visualization. A.R., T.T., M.M., M.B., N.W. wrote the manuscript.

## Notes

### Competing Interest Statement

The published work is actively used in Celeris Therapeutics GmbH to provide its services, and All authors are employees of said organisation

